# Individual differences in stereotypy and neuron subtype translatome with TrkB deletion

**DOI:** 10.1101/640987

**Authors:** Michel Engeln, Yang Song, Ramesh Chandra, Ashley La, Brianna Evans, Megan E. Fox, Shavin Thomas, T. Chase Francis, Ronna Hertzano, Mary Kay Lobo

**Author notes:** Corresponding author: Dr. Mary Kay Lobo, PhD, Department of Anatomy and Neurobiology, University of Maryland School of Medicine, 20 Penn Street, HSFII Rm 265, Baltimore, MD 21201, USA.

## Abstract

Motor stereotypies occurring in early-onset neuropsychiatric diseases are associated with dysregulated basal ganglia direct-pathway activity. Disruptions in network connectivity through impaired neuronal structure have been implicated in both rodents and humans. However, the neurobiological mechanisms leading to direct-pathway neuron disconnectivity in stereotypy remain poorly understood. We have a mouse line with Tropomyosin receptor kinase B (TrkB) receptor deletion from D1-expressing cells (D1-Cre-flTrkB) in which a subset of animals shows repetitive rotations and head tics with juvenile onset. Here we demonstrate these behaviors may be associated with abnormal direct-pathway activity by reducing rotations using chemogenetic inhibition of dorsal striatum D1-medium spiny neurons (D1-MSNs) in both juvenile and young adult mice. Taking advantage of phenotypical differences in animals with similar genotype, we then interrogated the D1-MSN specific translatome associated with repetitive behavior by using RNA-sequencing of ribosome-associated mRNA. Detailed translatome analysis followed by multiplexed gene expression assessment revealed profound alterations in neuronal projection and synaptic structure related genes in stereotypy mice. Examination of neuronal morphology demonstrated dendritic atrophy and dendritic spine loss in dorsal striatum D1-MSNs from mice with repetitive behavior. Together, our results uncover phenotype-specific molecular alterations in D1-MSNs that relate to morphological adaptations in mice displaying stereotypy behavior.

## Introduction

Stereotypies are repetitive, rhythmic behaviors ranging from simple movement sequences to complex actions^1^ that are observed in multiple neuropsychiatric diseases associated with basal ganglia dysfunction^1-3^. Imbalance in basal ganglia output-pathways arising from dopamine 1 receptor (D1, direct-pathway) vs. dopamine 2 receptor (D2, indirect-pathway) containing striatal medium spiny neurons (MSNs, aka-spiny projection neurons) is implicated in the repetitive movements that characterize disorders including Tourette Syndrome (TS), Autism Spectrum Disorders (ASD), and Obsessive Compulsive Disorder (OCD)^1, 3-6^. Activation of striatal dopamine D1 receptors induces stereotypies in non-human animals^7^, including those with dopamine deficiency^8^ and hyperdopaminergic function^9^, implicating a dysregulated direct-pathway in stereotypy behavior. Similarly, TS and OCD patients display increased sensitivity of striatal dopamine receptors and downregulation of D1 receptors^10, 11^ likely contributing to striatal hyperactivity^6, 12^. Additionally, mutations in genes regulating synaptic function and dendritic growth occur in TS, OCD and ASD^13-17^ and have been confirmed by striatal transcriptome analyses in TS and OCD patients^18, 19^. Thus, impaired MSN morphology combined with disrupted physiological properties may underlie neuronal hyperactivity and/or dopamine hypersensitivity and ultimately altered direct pathway output^8^.

Typically, stereotypies emerge during childhood and/or adolescence^3, 20^, suggesting a neurodevelopmental basis^21^. Striatal MSN development requires cortical brain-derived neurotrophic factor (BDNF) ^22, 23^. In humans, impairment of BDNF-Tropomyosin receptor kinase B (TrkB) signaling is associated with stereotypy disorders^24-26^ and conditional knockout of the BDNF-TrkB signaling pathway in striatal circuits leads to motor dysfunction including hyperlocomotion and stereotypy in mice^23, 27^. However, corticostriatal BDNF-TrkB knockout in mice usually affects both direct and indirect pathway MSNs to a similar extent, and BDNF deletion drastically impairs MSN development and survival^23, 27, 28^. In these conditions, parsing out the contribution of the BDNF-TrkB signaling in each pathway is complex. Cell-type specific TrkB deletion from enkephalin-expressing cells (D2-MSNs) induces spontaneous hyperlocomotion in middle-aged mice, which resembles Huntington’s disease (HD)^29^ and HD mouse models show impaired TrkB signaling in D2-MSNs^30^. Surprisingly, while evidence suggests that BDNF-TrkB signaling in direct pathway neurons could have an important role in the emergence of stereotypy, to date no study has assessed the effect of TrkB deletion in dopamine D1-MSNs on stereotypy behavior.

We have a mouse line in which the TrkB receptor is deleted from D1 expressing cells (D1-Cre-flTrkB). Interestingly, a subset of the animals (14.4%) exhibit repetitive circling behavior and head tics. Chemogenetic inhibition of striatal D1-MSNs reduced repetitive circling in the stereotypy mice. Thus, we took advantage of this phenotypic difference to probe molecular alterations associated with stereotypy in striatal D1-MSNs. Cell-type specific RNA-sequencing of ribosome-associated mRNA identified translatome alterations of morphology-related genes and morphological analysis demonstrated D1-MSN dendritic atrophy in mice with stereotypy.

## Materials and methods

### Experimental subjects

D1-Cre-flTrkB mice were generated by breeding D1-Cre (Line FK150)^31^ on a C57Bl/6 background with flTrkB^32^ mice on a mixed C57Bl/6 x FVB/N background. D1-Cre-TrkB^flox/+^ 33were crossed with TrkB^flox/+^ to obtain D1-Cre-TrkB^flox/flox^, D1-Cre-TrkB^flox/+^, and D1-Cre-TrkB^+/+^. For behavioral and neuronal morphology experiments, D1-Cre-TrkB^+/+^ (referred to as D1-Cre) and D1-Cre-flTrkB^flox/flox^ (referred to as D1-Cre-flTrkB) were used. TrkB knockout from D1-positive cells resulted in full-length TrkB deletion as described previously^32^. For cell-type specific RNA-sequencing studies D1-Cre-flTrkB mice were bred with RiboTag (RT)^+/+^ mice (Rpl22^tm1.1Psam/J^)^34^ on a C57Bl/6 background to generate D1-Cre-RT^+/-^ and D1-Cre-flTrkB-RT^+/-^ mice. No differences were observed between sexes in circling behavior: both males (26/55; i.e. 47.3%) and females (29/55; i.e. 52.7%) (**Supplementary Figure S1a**) were used in behavioral and neuronal morphology experiments. Studies were conducted in accordance with guidelines set up by the IACUC at University of Maryland School of Medicine (UMSOM). Animals were given food and water *ad libitum* and housed in UMSOM facilities during the whole study.

### Stereotypic behavior

Behavioral characterization was performed in 15 min weekly sessions between the age of 3 and 8 weeks in a 5L cylinder. Mice were first habituated to the cylinder for 15 min. In each session, rotations, head tics and grooming were video recorded and then analyzed. Rotations were defined as complete 360° turns^35^ while head tics were defined as uncontrolled head waving/shaking. The number of rearings was counted with video tracking software (CleverSys, Reston, VA, USA). For chemogenetic inhibition of D1-MSNs, a similar procedure was used. Briefly, mice were habituated to the cylinder before being injected with saline solution (i.p.;10mL/kg) 30 min prior to behavioral testing. On the next day, mice were injected with 0.5mg/kg clozapine-N-oxide (i.p.; LKT Laboratories, St. Paul, MN, USA) 30 min prior to behavioral testing.

### Stereotaxic surgery

Under isoflurane anesthesia, the dorsal striatum (from Bregma, anterior/posterior: +0.6, medial/lateral: +/-1.8, dorsal/ventral: −3) of juvenile (PND18-20) D1-Cre-flTrkB mice was bilaterally injected with a Cre-inducible, double inverted open (DIO)-reading frame adeno-associated virus (AAV): AAV5-hSyn-DIO-hM4D(Gi)-mCherry (Addgene; #44362) or AAV5-Ef1a-DIO-Cherry (UNC viral vector core) for Designer Receptor Exclusively Activated by Designer Drugs (DREADDs) experiments and with AAV5-DIO-eYFP (UNC vector core) diluted to 1.5×10^11^ VP/mL for neuronal morphology^36^. A minimum of 15 days was allowed for viral expression.

### Cell-type specific RNA sequencing and Bioinformatics

Immunoprecipitated polyribosomes were prepared from dorsal striatum of D1-Cre-RT and D1-Cre-flTrkB-RT mice according to our previous studies^36-38^. Briefly, four 12-gauge dorsal striatum punches were collected from a single animal and homogenized by douncing in homogenization buffer and 800 mL of the supernatant was added directly to the HA-coupled magnetic beads (Invitrogen: 100.03D; Covance: MMS-101R) for constant rotation overnight at 4°C. The following day, magnetic beads were washed three times in a magnetic rack for 5 min in high salt buffer. Finally, RNA was extracted with the RNeasy Micro kit (Qiagen: 74004) by adding TRK lysis buffer and following manufacturer’s instructions. For RNA-sequencing, RNA was quantified with a Bioanalyzer 2100 (Agilent) and only samples with RNA integrity numbers > 8 were used. Samples were submitted in biological triplicates for RNA-sequencing at the UMSOM Institute for Genome Sciences (IGS) and processed as described previously^39^. Libraries were prepared from 90ng of RNA from each sample using the NEBNext Ultra kit (New England BioLabs) per manufacturer’s instructions. Samples were sequenced on an Illumina HiSeq 4000 with a 75 bp paired-end read. An average of 66-97 millions reads were obtained for each sample. Reads were aligned to the mouse genome (Mus_musculus.GRCm38) using TopHat^40^ (version 2.0.8; maximum number of mismatches = 2; segment length = 30; maximum multi-hits per read = 25; maximum intron length = 50000) and the number of reads that aligned to the predicted coding regions were determined using HTSeq^41^. Significant differential expression (SDE) was assessed using DEseq. Post analysis was conducted to avoid over-inflation of fold change. In detail, all the values are quantile normalized Transcripts Per Kilobase Million (TPM) and values lower than 10% of the value of the dataset were replaced with the 10th quantile. Only genes with cutoff >1.2 (min(group1)/max(group2)>1.2, or min(group2)/max(group1)>1.2) and the absolute value of Log Fold Change (LFC) ≥ 1 in either of the pairwise comparisons were considered as differentially expressed (DE; 816 genes). All the DE genes in either of the pair-wise comparisons were uploaded into EXPANDER 2.0^42^ for non-supervised clustering using CLICK algorithm^43^ to find genes with similar patterns of expression and TANGO for GO functional enrichment analysis. Transcriptional regulator network were obtain with iRegulon^44^ by analyzing genes for GO terms based on CLICK cluster separation. Interaction maps between transcription factors and target genes were made using Cytoscape software^45^ (v. 3.6.1).

### RiboTag validation and Gene expression

To validate the specific enrichment of D1-MSN associated genes (*Drd1, Chrm4, Tac1*), we synthetized 100ng of cDNA from immunoprecipitated and isolated polyribosome-associated RNA (see above for detailed procedure) as well as non-precipitated RNA (input) using the iScript cDNA synthesis kit (Bio-Rad, USA). cDNA was then preamplified using TaqMan assay following kit instructions^46^. FAM probes for genes of interest (20X TaqMan Gene Expression Assay, Applied Biosystems; see **Supplementary Table 1** for primer information) were multiplexed and diluted to a final concentration of 0.2X and amplified for 14 cycles using a PCR thermal cycler (C1000 Touch, Biorad). After amplification, the samples were diluted at a 1:10 concentration with TE buffer. qRT-PCR was run with TaqMan Gene expression Master Mix (2X, Applied Biosystems) on a CFX-384 Touch (Bio-Rad, USA) and quantification of mRNA changes was performed using the ΔΔC_T_ method^47^.

RNA-sequencing data were confirmed in a separate set of RiboTag processed samples. Following polyribosome immunoprecipitation and RNA isolation, cDNA was synthetized with iScript cDNA synthesis kit (Bio-Rad, USA). 10ng of cDNA was amplified using the Low Input Kit (NanoString technologies, USA) before being processed with nCounter Master kit according to manufacturer instruction (NanoString Technologies, USA) by UMSOM IGS on a custom-made gene expression Code set. Primer sequences for NanoString are available in **Supplementary Table 1**. Data were analyzed with nSolver Analysis software as in^48^.

### Immunostaining

Mice were perfused with 0.1M PBS and 4% paraformaldehyde. Brains were post-fixed for 24 hours and sections were collected in PBS with a vibratome (Leica, Germany). For DREADDs virus validation, 40µm sections were blocked in 3% normal donkey serum (NDS) and 0.3% Triton X-100 in PBS for 30 min, then incubated in rabbit anti-ds-Red (1:1500; Clontech: #632496) overnight at room temperature. Slices were washed 3 × 10 min in PBS prior to 2 hours incubation in anti-rabbit-Cy3 (1:1000, Jackson Immuno; #111-166-003). Slices were washed in PBS, mounted with Vectashield containing DAPI, and imaged on a confocal microscope. For D1-MSN morphology, 100 µm slices were washed 3 × 5 min with PBS and blocked as described above. Slices were incubated in chicken anti-GFP (1:500; Aves Lab, #GFP-1020) at 4°C overnight. Slices were washed 3 × 5 min, then 7 × 60 min, then incubated in Anti-Chicken Alexa 488 (1:500; Jackson Immuno, #111-545-144) at 4°C overnight. Slices were washed with PBS and mounted with Vectashield mounting media^36^.

### Neuronal morphology

Sections containing dorsal striatum from 8-week-old mice were sampled from bregma AP: 1.2-0.6 mm and Z-stack images were acquired on a confocal microscope (Olympus FV500) at 0.6 µm increments using a 40× objective. As described previously^36, 37^, D1-MSNs were 3D reconstructed using Imaris 8.3 software (Bitplane, Oxford Instruments). Surfaces were masked to generate a 2D image of a single MSN for sholl analysis. Concentric ring intersections were determined using the ImageJ sholl analysis plugin^49^ at 10 µm increments from soma. Spine images were acquired from secondary dendrites with 0.2 µm increments Z-stacks using a 60x objective with 2x digital magnification and quantified with Neuron Studio software^50^. For all morphological analyses, 3-4 cells were averaged per animal.

### Statistical analysis

Graphpad Prism 6.0 software was used for statistical analysis. Normality of the data was assessed with Bartlett’s test. ANOVAs and Kruskal-Wallis tests were run for normal and non-normal data respectively. Tukey, Sidak and Dunn’s post hoc tests were used. Samples were excluded if not detected (molecular analysis) or if they failed Grubbs’ outlier test. Sample sizes were determined from previous studies using mouse behavior, cell-type specific RNA isolation and neuronal morphology^36-38^. In figure legends: *p<0.05, **p<0.01, ***p<0.001. All graphs represent mean ± s.e.m. In graphs, individual values are plotted to report that variation and variance is similar between groups that are compared.

## Results

### A subset of D1-Cre-flTrkB mice exhibit D1-MSN related repetitive behavior

While the majority of D1-Cre-flTrkB mice showed no stereotypy (further referred to as D1-Cre-flTrkB-NS), behavioral characterization revealed that a subset of both male and female D1-Cre-flTrkB mice (further referred to as D1-Cre-flTrkB-S) exhibited stereotypic behavior (55/381; 14.4%; **Supplementary Figure S1a, Supplementary Video 1**). D1-Cre-flTrkB-S displayed significantly more rotational behavior: 2-Way RM-ANOVA: Group: F_(2;29)_= 21.42; p<0.0001; Tukey post-hoc: p<0.05 when compared to D1-Cre and D1-Cre-flTrkB-NS from the ages 3 to 8 weeks (**Figure 1a**). The animals lacked a preference towards one side since both clockwise and counter-clockwise circling was observed. D1-Cre-flTrkB-S was the only group to exhibit head tics during that same time period: 2-Way RM-ANOVA: Group: F_(2;29)_= 63.83; p<0.0001; Tukey post-hoc: p<0.01 (**Figure 1b**). No differences in rearing behavior were observed between groups: 2-Way RM-ANOVA: Group: F_(2;29)_= 1.198; p=0.32 (**Supplementary Figure S1b**) and although knockout mice from both phenotypes exhibited lower grooming time at 3 and 4 weeks of age, this difference did not persist at later ages: 2-Way RM-ANOVA: Group: F_(2;29)_= 6.26; p<0.01, Tukey post-hoc: p<0.05 for 3 and 4 weeks and p>0.05 from 5 to 8 weeks (**Supplementary Figure S1c**). Balanced activation between direct and indirect pathways controls movement^51^. To see if the stereotypy is mediated through striatal D1-MSNs we then used inhibitory chemogenetics in D1-MSNs. We infused a Cre-dependent hM4D(Gi) inhibitory DREADDs in the dorsal striatum to transiently inhibit D1-MSN synaptic transmission^52^ (**Figure 1c; Supplementary Figure S1d**). Inhibition of D1-positive cells can drastically decrease locomotor activity^53^ and we wanted to reduce repetitive behavior without altering normal movement. Low concentrations of CNO can partially decrease synaptic transmission in intact neuronal circuits^52^. We thus used a 0.5 mg/kg (i.p.) clozapine-N-oxide (CNO) dose that had no effect on locomotion in an open field (**Supplementary Figure S1e**) and that was insufficient to impact rotational behavior in the absence of the hM4D(Gi) receptor (**Supplementary Figure S1f**). In humans, stereotypic behaviors appear during childhood and persist through adolescence^3, 54^. We thus tested, in two separate cohorts, the effect of D1-MSN inhibition at both juvenile (4 weeks) and young adult (8 weeks) ages. D1-Cre-flTrkB-S mice receiving CNO displayed a significant 27% decrease in circling behavior relative to pre-CNO vehicle injection at 4 weeks: 2-Way RM-ANOVA: interaction Group x Treatment: F_(2,21)_= 4.87; p= 0.02; Sidak post-hoc: p<0.01 (**Figure 1d**) and a significant 31% decrease relative to pre-CNO vehicle injection at 8 weeks: 2-Way RM-ANOVA: interaction Group x Treatment: F_(2,20)_= 8.71; p= 0.002; Sidak post-hoc: p<0.001 (**Figure 1e**). However, CNO had no effect on head tics at both 4 and 8 weeks of age: paired t-test: t=0.37, df=7; p= 0.72 and t=0.37, df=6; p= 0.22 respectively (**Supplementary Figure S1g**). In both D1-Cre and D1-Cre-flTrkB-NS mice, D1-MSN inhibition had no effect on rotations further confirming that our inhibition protocol had no impact on normal motor behavior (**Figure 1d, e**).

**Figure 1:**
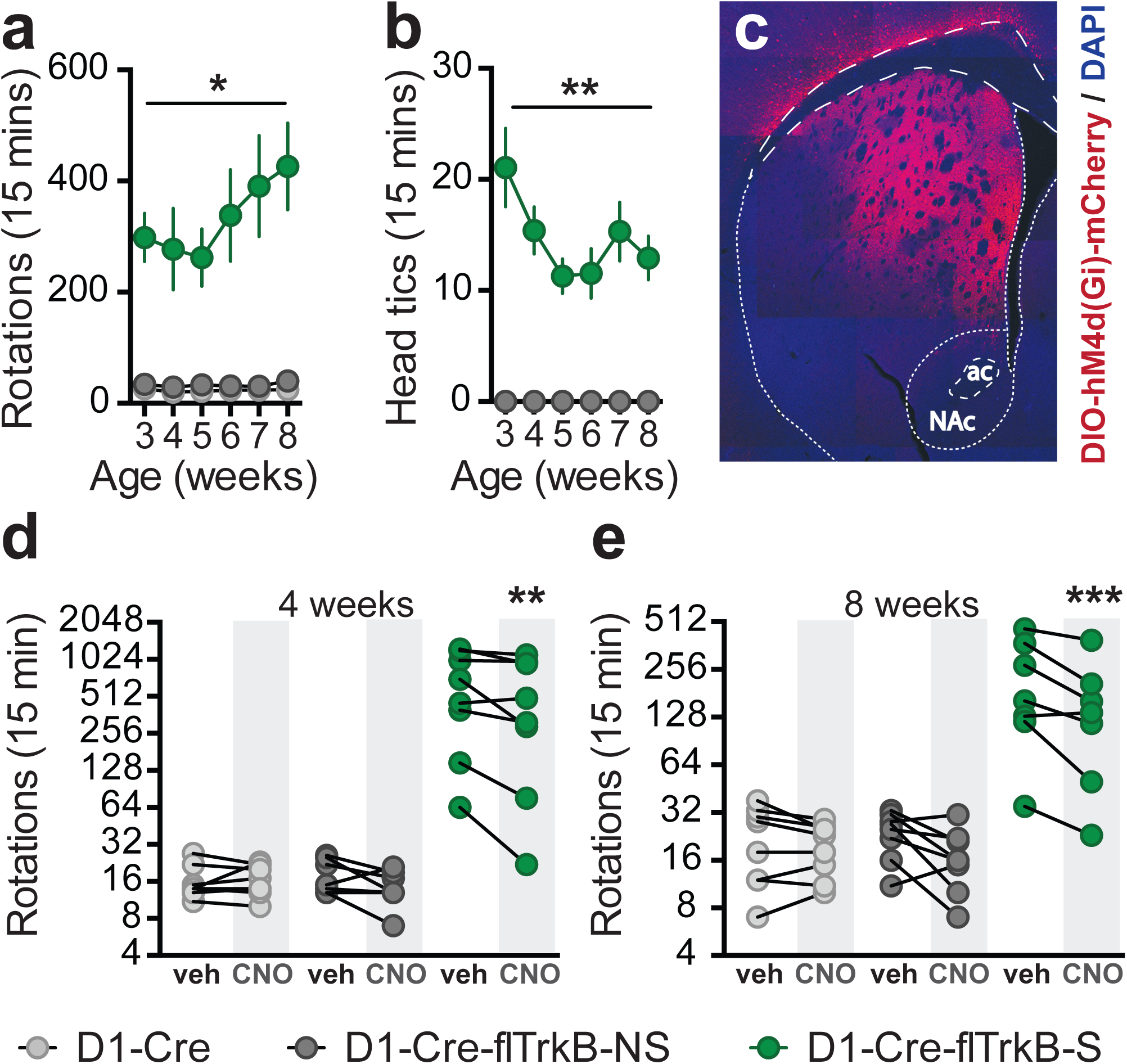
Behavioral characterization of D1-Cre-flTrkB mice: **a.** Rotational behavior in mice displaying D1-Cre-flTrkB-S, D1-Cre-flTrkB-NS, and D1-Cre control mice from juvenile to young adult ages (*p<0.05); **b.** Only D1-Cre-flTrkB-S mice display head tics between the age of 3 and 8 weeks (**p<0.01); (a-b) n= 13 D1-Cre-flTrkB-S, 7 D1-Cre-flTrkB-NS, and 12 D1-Cre; **c.** Representative image of AAV-DIO-hM4d(Gi)-mCherry virus injection in the dorsal striatum (ac= anterior commissure, NAc= nucleus accumbens); **d.** Rotational behavior following vehicle (veh) and 0.5 mg/kg (ip.) clozapine-N-oxide (CNO) in 4 week-old mice (**p<0.01), n=8 in each group; **e.** Rotational behavior following vehicle and 0.5 mg/kg (ip.) CNO in 8 week-old mice (***p<0.001), n= 8, 8, 7 respectively. For ‘Rotations’, please note the scale of the y-axis.

### Molecular profiling of striatal D1-MSNs

First, using the RiboTag procedure we demonstrate significant enrichment of D1-MSN-related mRNAs in immunoprecipitated sample relative to input from dorsal striatum (t-test: Drd1: t=4.22; Chrm4: t=2.36; Tac1: t=3.03; df=18, p<0.05; **Supplementary Figure S2a**), consistent with our previous studies in ventral striatum using this methodology and consistent with other studies in dorsal striatum ^36-38, 47, 55-57^. Since individual differences in stereotypy behavior were observed between D1-Cre-flTrkB-S vs. D1-Cre-flTrkB-NS mice and stereotypy was partially reduced by chemogenetic inhibition of striatal D1-MSNs, we then probed molecular adaptations in the D1-MSNs. We used D1-Cre-flTrkB-RiboTag (RT)- S, -NS, and D1-Cre-RT control mice, at age 4 weeks when stereotypy behavior is observed and reduced by chemogenetic inhibition of striatal D1-MSNs. Ribosome-associated mRNA was isolated from dorsal striatum D1-MSNs from each group followed by translatome profiling (**Figure 2a**). Using RNA-sequencing, we identified a total of 14,764 genes expressed in our samples with fold of change greater than 1 in either of the pair-wise comparisons. **(Supplementary Figure S2b; Supplementary Table 2)**.

**Figure 2:**
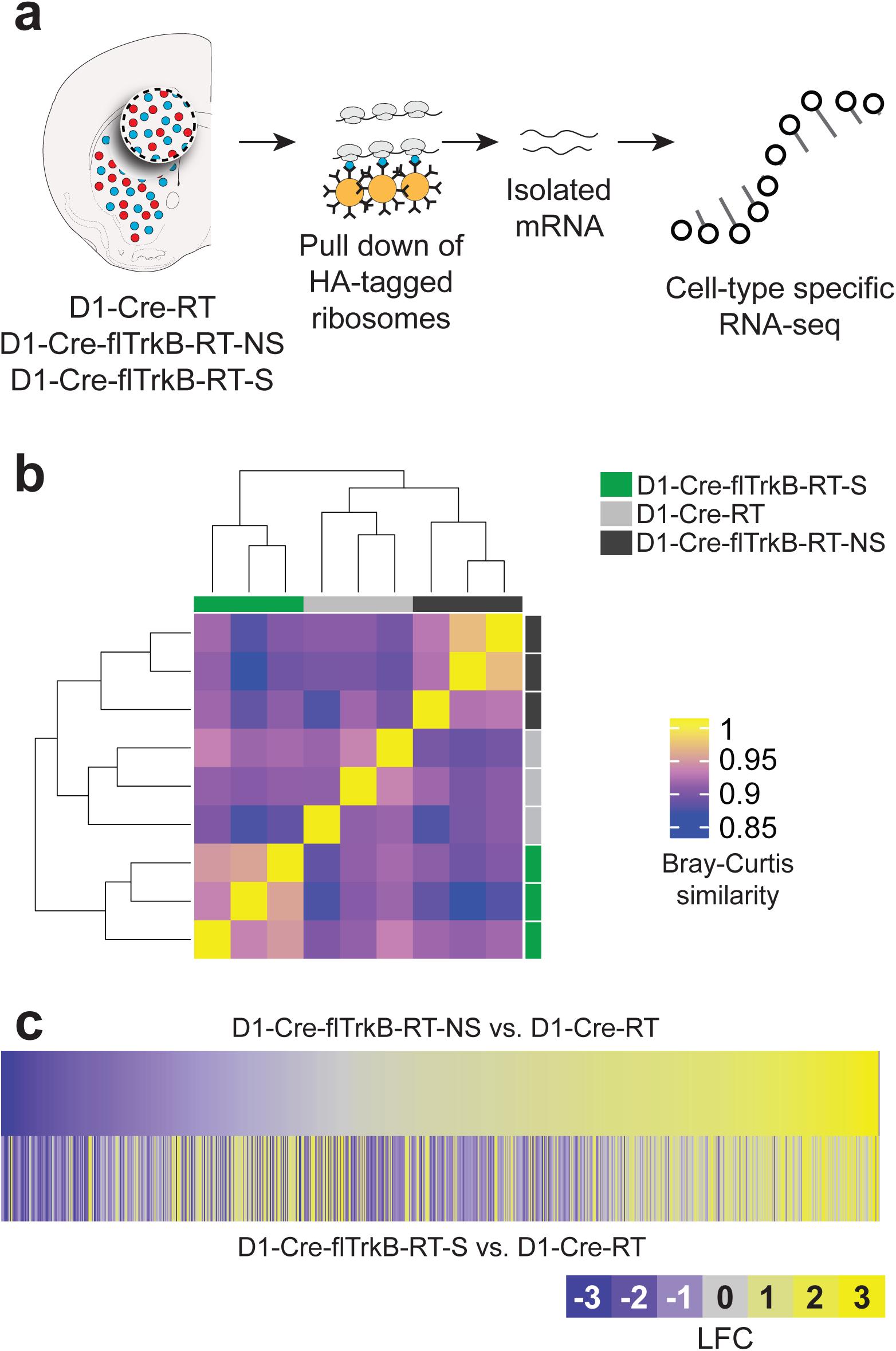
D1-MSNs of mice with stereotypy display distinct translatome profiles: **a.** Schematic of the RiboTag isolation of D1-MSN mRNA followed by cell-type specific RNA-sequencing; **b.** Heat map and dendrogram for unbiased hierarchical clustering (scale= Bray-Curtis similarity values); **c.** Heatmaps of the 816 expressed genes displaying differential gene expression for D1-Cre-flTrkB-RT-NS and D1-Cre-flTrkB-RT-S mice compared to D1-Cre-RT (LFC= Log fold change).

Interestingly, the overall translatome profile of D1-MSNs in D1-Cre-flTrkB-RT-NS mice was more similar to D1-Cre-RT than to D1-Cre-flTrkB-RT-S mice (**Figure 2b**). However, differences in gene expression between D1-Cre-flTrkB-RT-NS and D1-Cre-flTrkB-RT-S emerged (**Supplementary Figure S2b**), which may reflect that repetitive behaviors lead to different impact on gene expression level. Overall, we found 816 genes with significantly different expression (**Figure 2c; Supplementary Table 3**). Based on non-supervised clustering algorithm (CLICK algorithm), all the differentially expressed genes were grouped to 4 clusters (overall average homogeneity= 0.807; overall average separation= −0.336): 1: lower expression in D1-Cre-flTrkB-RT-S mice (338 genes); 2: higher expression in D1-Cre-flTrkB-RT-S (245 genes); 3: lower expression in D1-Cre-RT (122 genes); 4: higher expression in D1-Cre-RT (111 genes) (**Figure 3a; Supplementary Table 3**). Ontology (GO) analysis was further conducted of these 4 clusters. Cluster 1 presented the most GO terms (i.e. 124) compared to Cluster 2 (53 terms), Cluster 3 (9 terms) and Cluster 4 (36 terms). Among the 124 enriched GO categories in Cluster 1, which corresponds to genes showing decreased expression in D1-Cre-flTrkB-RT-S, 43 were related to neuronal, dendritic and synaptic development as well as receptor and/or transporter activity and signaling. Of particular interest, the GO term ‘Regulation of signal transduction’ (GO:0009966; 74 genes; 22% frequency in set; p= 2.42 E-14), ‘Cell projection organization’ (GO:0030030; 34 genes; 10% frequency in set; p= 6.62 E-9), ‘Cell projection morphogenesis’ (GO:0048858; 27 genes; 8% frequency in set; p= 6.10 E-13), ‘Regulation of neuron projection development’ (GO:0010975; 23 genes; 6.8% frequency in set; p= 3.02 E-8), ‘Regulation of synapse structure or activity’ (GO:0050803; 12 genes; 3.6% frequency in set; p= 9.82 E-8) were significantly changed in D1-Cre-flTrkB-RT-S mice (**Figure 3b**). In Cluster 2, which corresponds to genes with increased expression in D1-Cre-flTrkB-RT-S, one GO term with significant enrichment particularly reflected the phenotype present in our stereotypy mice: ‘Regulation of locomotion’ (GO:004001227; 29 genes; 12% frequency in set; p= 1.78 E-12; **Figure 3b**). Detailed analysis of gene expression for ‘Cell projection morphogenesis’ (GO:0048858) revealed that, out of 27 genes showing differential expression in our dataset, 11 were significantly different in D1-Cre-flTrkB-RT-S compared to D1-Cre-RT and 25 compared to D1-Cre-flTrkB-RT-NS (**Figure 3c; Supplementary Table 3**). Similarly for the other GO terms identified here, D1-Cre-flTrkB-RT-S showed significant differences for 82 to 94% of genes from Cluster 1 and for 62% of genes from Cluster 2 when compared to D1-Cre-flTrkB-RT-NS (**Supplementary Figure S2c; Supplementary Table 3**). Together, the GO analysis suggested that D1-MSNs from mice with stereotypy display altered gene expression for molecules in dendritic and synaptic pathways.

**Figure 3:**
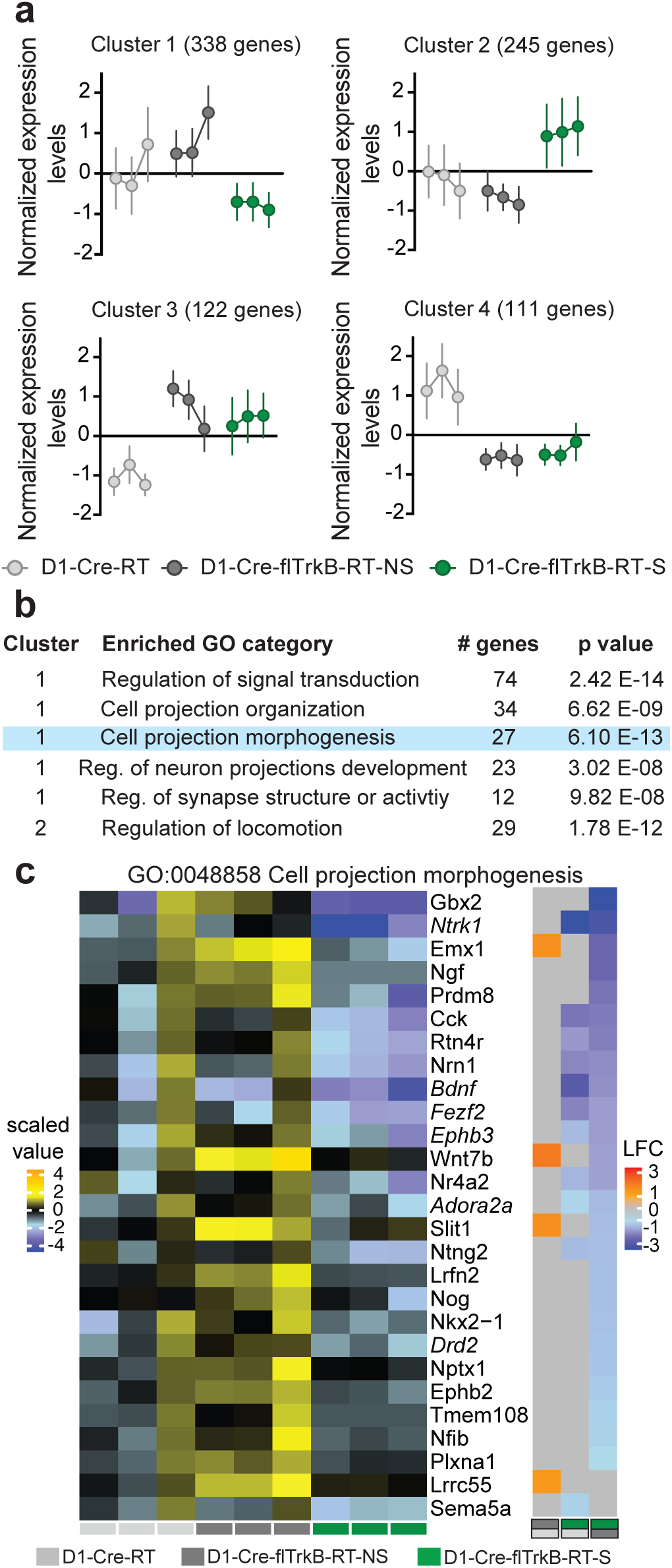
Gene ontology analysis reveals alterations in morphology-related genes in D1-MSNs of mice with stereotypy: **a.** Non-supervised clustering algorithm (CLICK algorithm) identified 4 clusters based on gene expression: 1: low expression (338 genes) and 2: high expression (245 genes) in mice with stereotypy, 3: low expression (122 genes) and 4: high expression (111 genes) in control mice; **b.** List of the top GO terms related to neuronal morphology and motor function. Blue highlight denotes the GO term detailed in the following panel; **c.** Left: Heat map for the biological replicates for the GO term “Cell projection morphogenesis”, Right: Pairwise comparisons for the corresponding genes in the 3 conditions: D1-Cre-flTrkB-RT-S showed decreased expression for 25 genes when compared to D1-Cre-flTrkB-RT-NS and 11 genes when compared to D1-Cre-RT mice (LFC= Log fold change). Light grey (D1-Cre-RT), dark grey (D1-Cre-flTrkB-RT-NS) and green (D1-Cre-flTrkB-RT-S) labels below the heat maps indicate groups and group comparisons. Genes in italic were selected for multiplexed mRNA measurements (see Figure 4).

To further identify whether transcriptional regulons are also impacted by stereotypy, we further conducted transcriptional regulator network analysis using iRegulon^44^. Three of the GO terms we identified here appeared to share a common transcriptional regulator. The transcription factor *Nr2f1* shares a binding motif with: 5 genes (42%) of the ‘Regulation of synapse structure or activity’ term: motif similarity: FDR< 8.136 E-4; 22 genes (30%) of the ‘Regulation of signal transduction’ term: motif similarity: FDR< 2.502 E-4; 19 genes (66%) of the ‘Regulation of locomotion’ term: motif similarity: FDR< 1.122 E-4 (**Figure 4a**). Based on this list as well as genes involved in ‘Cell projection morphogenesis’ (**Figure 3c**), we selected several target genes and measured multiplexed mRNA levels in D1-MSNs at both 4 and 8 weeks using NanoString. The expression of a subset of these genes was not detected at 4 weeks but reached detection threshold at 8 weeks. Additionally, many genes showed trends of altered expression at 4 weeks and reached significance at 8 weeks (**Figure 4b**). To account for this evolution in gene expression, we reported both significant (p<0.05) and trending (p≤0.09) results. At 4 weeks, the expression of genes such as *Macf1, Mme, Ndnf, S1pr1* and *Strn* was altered in D1-Cre-flTrkB-RT-S mice (**Figure 4b, Supplementary Table 4**). Alterations in gene expression continued at 8 weeks for over 60% of the genes from our list in D1-Cre-flTrkB-RT-S mice. Overall, most genes showed decreased levels, which is consistent with Cluster 1 profile, and genes associated with ‘Regulation of locomotion’ were upregulated following Cluster 2 profile (**Figure 4b; Supplementary Table 4**). Interestingly the expression of *Nr2f1*, which regulates the expression of numerous genes in our list, was significantly reduced in D1-Cre-flTrkB-RT-S mice at 8 weeks. Finally at both time points, the expression of *Ntrk2* (TrkB) was drastically decreased in both D1-Cre-flTrkB-RT-NS and D1-Cre-flTrkB-RT-S mice confirming both the effective knockout of TrkB in our mice as well as the cell-type specificity of our RiboTag samples (**Figure 4b; Supplementary Table 4**). Of note, we also observed distinct patterns of expression of D2-MSN enriched genes: *Adora2a, Drd2* and *Penk* mRNA was upregulated in non-stereotypy mice and down-regulated in stereotypy mice at 8 weeks (**Figure 4b; Supplementary Table 4**). *Penk* was also reduced in stereotypy mice at 4 weeks. While D1-MSN enriched genes did not display distinct patterns of expression, we did observe a significant change in two of these genes: *Chrm4* was upregulated in D1-Cre-flTrkB-S at 4 weeks and *Pdyn* was upregulated in D1-Cre-fltrkB-NS at 8 weeks (**Figure 4b; Supplementary Table 4**).

**Figure 4.**
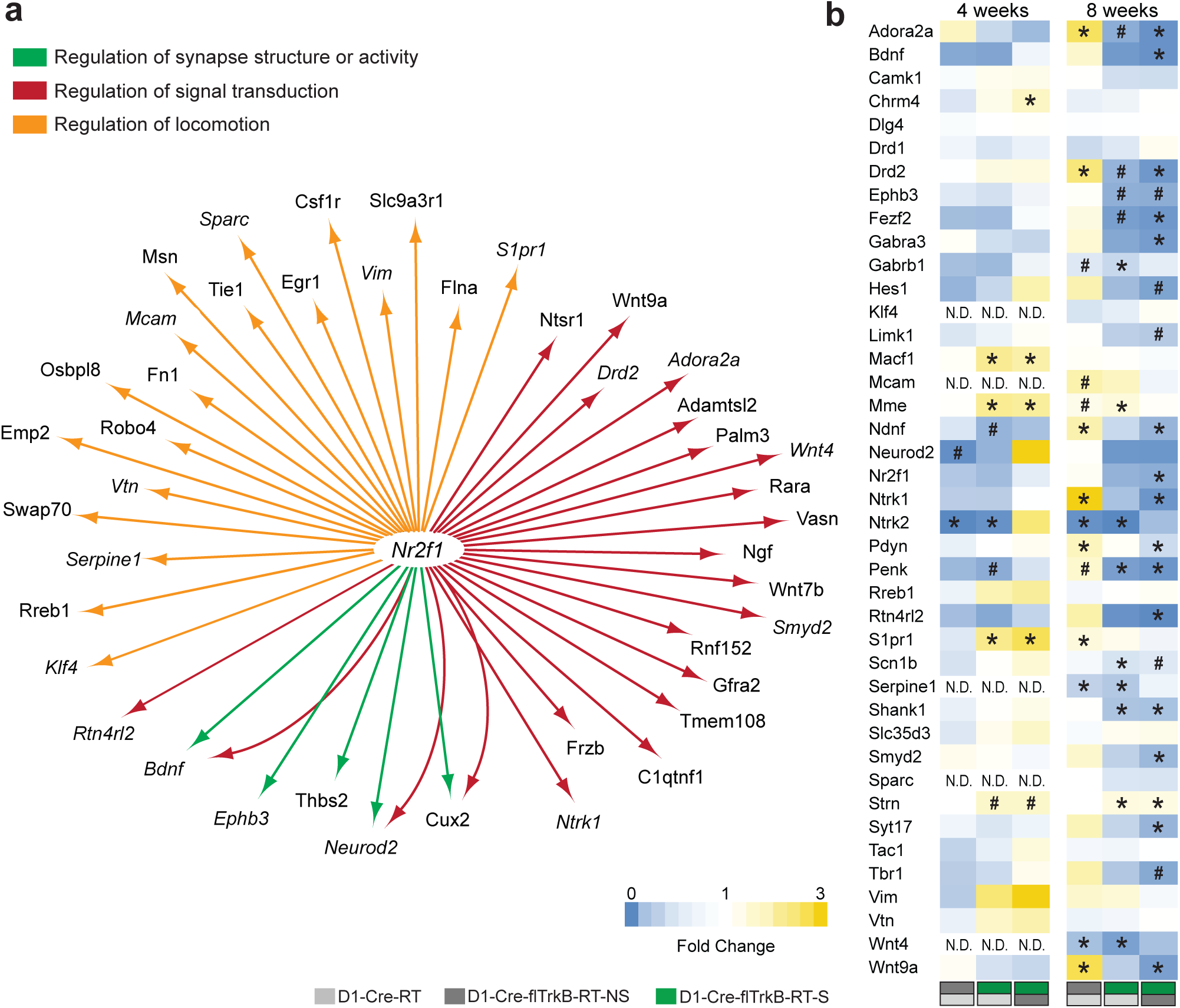
Multiplexed mRNA analysis of D1-MSNs. **a.** Transcriptional regulator network analysis shows that *Nr2f1* is a common regulator for multiple morphology-related genes. Genes in italic were selected for multiplexed mRNA measurements; **b.** Multiplexed mRNA analysis (NanoString) on both juvenile and young adult mice (# p≤0.09, *p<0.05). Light grey (D1-Cre-RT), dark grey (D1-Cre-flTrkB-RT-NS) and green (D1-Cre-flTrkB-RT-S) labels below the heat maps indicate groups and group comparisons.

### Stereotypy mice exhibit D1-MSN atrophy

The translatome analyses pointed toward morphological alterations in D1-MSNs of mice with stereotypy behavior. With most translatome alterations related to neuronal morphology displaying significance at 8 weeks of age, we decided to evaluate neuronal morphology at this age. We sparsely labeled dorsal striatum D1-MSNs with AAV-DIO-eYFP and assessed different morphological parameters (**Figure 5a**). First, we observed a decreased number of branch points: 1-way ANOVA: F_(2;12)_= 5.31; p= 0.02; Tukey post-hoc: p<0.05 (**Figure 5b**) and a reduction in the complexity of D1-MSNs from D1-Cre-flTrkB-S mice as shown by reduced Sholl intersections: 2-way RM-ANOVA Group: F_(2;252)_= 29.65; p< 0.0001; Tukey post-hoc: p<0.05 (**Figure 5c**). Similarly, we found that D1-Cre-flTrkB-S mice have significantly shorter dendrites: 1-way ANOVA: F_(2;12)_= 4.48; p= 0.03; Tukey post-hoc: p<0.05 (**Figure 5d**). However, soma volume was unchanged in these mice (**Figure 5e**), suggesting that morphological alterations were not due to general neuronal reduction. Thus, as indicated by the terms ‘Cell projection organization’, ‘Cell projection morphogenesis’ and ‘Regulation of neuron projection development’ we confirmed that D1-Cre-flTrkB-S mice displayed altered D1-MSN morphology. Additionally, since the GO analysis suggested altered synaptic structure: ‘Regulation of synapse structure or activity’ and ‘Regulation of signal transduction’, we measured dendritic spines in all three groups (**Figure 5f**). Spine density was significantly reduced in D1-Cre-flTrkB-S mice: 1-way ANOVA: F_(2;12)_= 5.07; p= 0.03; Tukey post-hoc: p<0.05 (**Figure 5g**).

**Figure 5:**
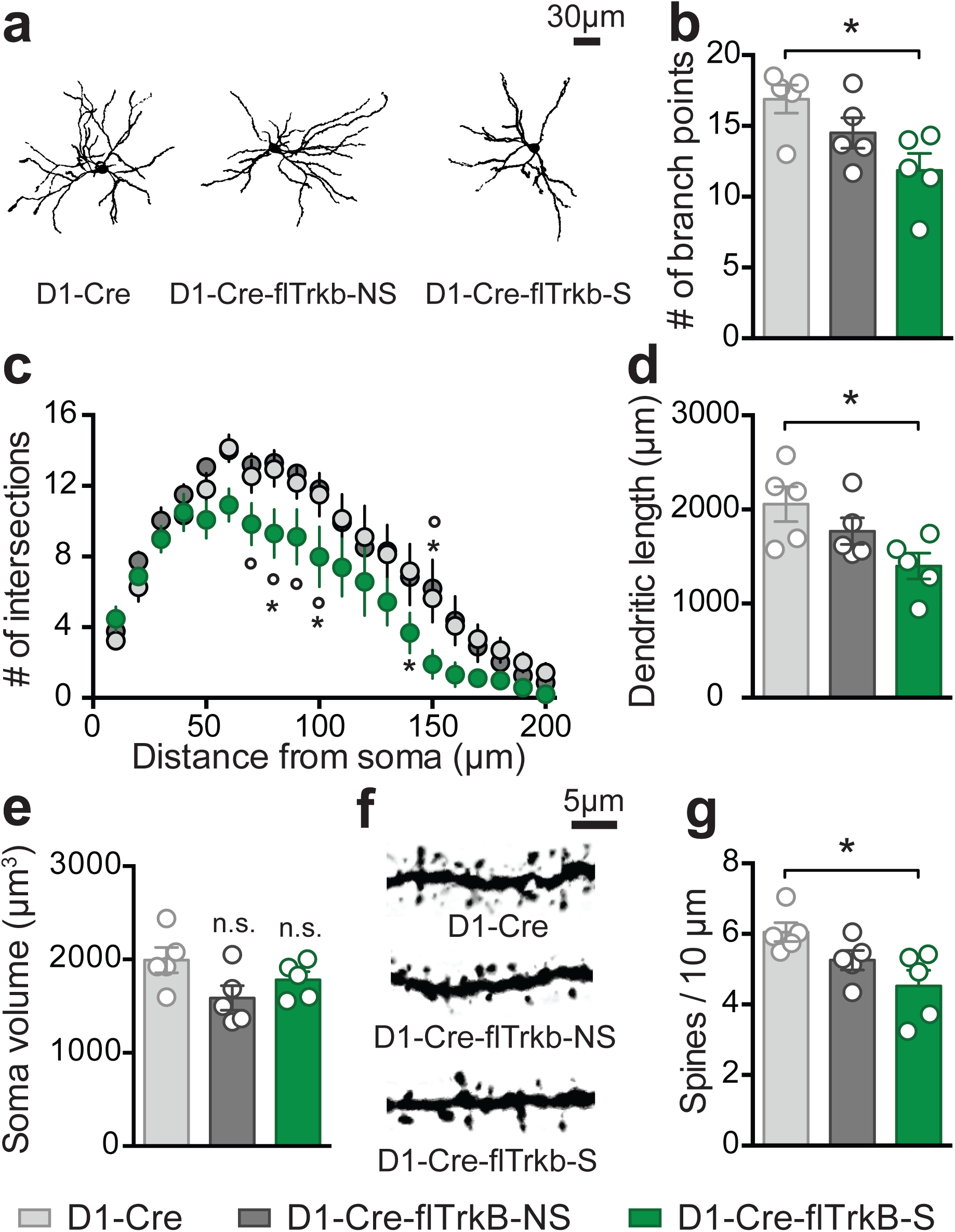
Mice with stereotypy display D1-MSN dendritic atrophy and reduced dendritic spines: **a.** Representative images of dorsal striatum D1-MSN from each group; **b.** Decreased neuronal complexity (Sholl analysis, *p<0.05 from D1-Cre and °p<0.05 from D1-Cre-flTrkB-NS); **c.** number of branch points (*p<0.05) and, **d.** dendritic length (*p<0.05) in D1-Cre-flTrkB-RT-S mice; **e.** Soma volume is unchanged in both D1-Cre-flTrkB-NS and D1-Cre-flTrkB-S mice; **f.** Representative images of D1-MSN dendritic spines from each group; **g.** Decreased spine density in D1-Cre-flTrkB-S mice (*p<0.05). For all graphs, n= 5 mice/group, 3-4 cells/mouse.

## Discussion

In this study we showed that TrkB knockout from D1-expressing cells results in stereotypic rotational behavior and head tics in a subset of mice. Reduced repetitive circling following chemogenetic inhibition of striatal D1-MSNs supports a role for these neurons in mediating the stereotypy behavior. By comparing the cell-type specific translatome in mice with a similar genotype but different phenotype, we showed that D1-MSNs from mice with stereotypy displayed alterations in ribosome-associated mRNAs for gene networks including those regulating neuronal morphology and locomotion. Finally, we observed dendritic atrophy and reduced dendritic spines in D1-MSNs of D1-Cre-flTrkB-S mice, which is consistent with alterations in neuronal morphology related gene networks.

Stereotypies are associated with increased direct motor pathway activity^1, 3, 8, 9^. In human patients, impairments in BDNF-TrkB signaling are associated with several stereotypy disorders^24-26^. Here, we found that only a subset of mice lacking the TrkB receptor in direct-pathway neurons exhibit repetitive circling behavior. Stereotypies appear during childhood and/or adolescence in humans^3, 20^. We thus interrogated the translatome of dorsal striatum D1-MSNs in juvenile mice using cell type-specific RNA-sequencing. First, unbiased hierarchical clustering performed on the entire dataset revealed that similarities in gene expression pattern were not shared by mice with similar genotype but rather by similar phenotype. This suggested that although D1-MSNs of D1-Cre-flTrkB mice undeniably share similarities in gene expression likely related to TrkB knockout (see Cluster 3 and 4 of **Figure 3a**), transcriptional alterations in D1-MSNs associated with repetitive behavior were greater than the effect of gene deletion alone. This was supported by the non-supervised clustering analysis (CLICK algorithm), revealing that 71% of the post-filtrated genes with significant changes belonged to the mice with stereotypy.

To obtain better insight into the alterations present in D1-MSNs of mice with stereotypy, we conducted a gene ontology analysis on Cluster 1 and 2 (i.e. down-or up-regulated genes in mice with repetitive behavior). Interestingly, 35% of the down-regulated genes were related to dendritic and synaptic development as well as receptor and/or transporter activity and signaling. Detailed analysis of genes belonging to these GO categories allowed us to generate a list of genes that can potentially support cellular adaptations underlying repetitive behavior as well as a common transcriptional regulator. Multiplexed mRNA analysis of 4-week-old animals confirmed key genes coding for cytoskeleton, neurite outgrowth, spines or synapse formation such as *Macf1*^58^, *Mme*^59^, *Ndnf*^60^, *Strn*^61^ or *S1pr1*^62^ had altered expression in mice with repetitive behavior. These genes have fundamental roles throughout brain development. Impaired expression, observed here at juvenile age, could support altered D1-MSN functioning leading to repetitive behaviors. While some transcriptional changes were already present at 4 weeks, more reached statistical significance at 8 weeks of age. This might reflect neural development processes in mice during which neurotransmission, synaptic function and neuronal networks undergo maturation and pruning stages between PND 25 and PND 49 to reach a stable, fully formed, adult brain at PND >60^63^. Additionally, the establishment of transcriptional differences at 8 weeks in mice with stereotypy could be due to a gradual decreased expression of *Nr2f1* (COUP-TF1), which was significantly decreased at 8 weeks. *Nr2f1* was identified as a potential common transcriptional regulator for multiple genes in our dataset and is involved in neurite outgrowth^64^. Impaired transcriptional regulation could thus have a broad impact on neuronal morphology. Altogether, our mRNA data confirmed a wide range of transcriptional alterations in stereotypy mice and suggested that differential expression of morphology-related genes persist until young-adult stages.

Based on these translatome findings, we evaluated D1-MSN morphology in young adult mice. In mice with stereotypy, we observed decreased dendritic complexity and dendritic length. Striatal neurons integrate cortical and subcortical information to provide error detection monitoring in order to generate harmonious motor behaviors^6, 65^. D1-MSN dendritic atrophy could result in altered functional connectivity and aberrant signal processing supporting abnormal behavior^66-68^. Reduced expression of the spine-related genes *Shank1* and *Strn*, both associated with impaired circuitry in ASD^61, 69^, as well as decreased spine density further suggest reduced cortical inputs onto MSNs. Additionally, reduced excitatory input can induce changes in intrinsic excitability at the postsynaptic level to maintain homeostasis in circuit activity^70^. Our group has recently described this mechanism for stress-induced dendritic atrophy in ventral striatal D1-MSNs^36, 38, 71^. Further, our previous studies demonstrate reduced inhibition of ventral striatal D1-MSNs in D1-Cre-flTrkB mice^72^. In mice with stereotypy, reduced dendritic complexity could thus lead to a homeostatic increase in excitability or reduction in inhibition in dorsal striatal D1-MSNs. This is consistent with chemogenetic inhibition of D1-MSNs reducing circling behavior in stereotypy mice. This is also in line with a decrease in the voltage-gated sodium channel subunit gene, *Scn1b*, in D1-MSNs of stereotypy mice, since loss of function of this molecule can lead to the development of hyperexcitability in the brain^73^.

Along with alterations in morphology and locomotor related molecules we observed changes in D2-MSN enriched genes: *Adora2a, Drd2*, and *Penk*. mRNA for these genes was significantly upregulated in D1-MSNs of non-stereotypy mice and downregulated in stereotypy mice at 8 weeks. *Penk* was also significantly reduced in stereotypy at 4 weeks. This could reflect a greater proportion of MSNs expressing both D1 and D2 enriched molecules in non-stereotypy mice. A previous study showed that enhancing *Drd2* expression in striatum leads to increased presence of D1-MSN bridging collaterals to the globus pallidus externus (GPe) and enhanced inhibition of GPe neurons^74^. A similar mechanism could be occurring here with enhanced *Drd2* in D1-MSNs of non-stereotypy mice. Given the morphological and physiological differences between both MSN subtypes^75, 76^, this could contribute to stabilized activity between direct and indirect pathways in these mice. While patterns of expression in groups of D1-MSN enriched genes was not observed between stereotypy and non-stereotypy mice we did observe an increase of the striosome D1-MSN enriched gene, *Pdyn*-the precursor for the dynorphin peptide^77^, in D1-MSNs of non-stereotypy mice at 8 weeks. Further, *Pdyn* was reduced in D1-MSNs of stereotypy mice when compared to non-stereotypy mice. In contrast mRNA for the matrix D1-MSN enriched gene, *Chrm4* ^78^ -the gene for acetylcholine muscarinic receptor 4, is enhanced in D1-MSNs of stereotypy mice at 4 weeks. The differential regulation of *Pdyn* vs. *Chrm4* could reflect an imbalance in striosome vs. matrix function, which is implicated in stereotypy behavior^2^. Additionally, the responsiveness of stereotypy mice to the low dose of CNO, acting on the DREADD hM4D(Gi), derived from *Chrm4*, could be a consequence of enhanced sensitivity of M4 signaling that might occur through upregulation of endogenous *Chrm4*.

In summary, our study reveals deletion of TrkB in D1 expressing neurons leads to individual differences in repetitive behavior, that are associated with altered D1-MSN translatome and dendritic morphology. These cellular alterations potentially disrupt D1-MSN output activity to promote repetitive behavior. How the differences in D1-MSN translatomes between D1-Cre-flTrkB mice with and without stereotypy emerge remains unclear and could involve point mutations associated with impaired BDNF-TrkB signaling^26^ that may either aggravate or compensate for the effect of TrkB knockout. Insight into the distinct molecular and cellular adaptations that underlie stereotypy or lack of stereotypy behavior could uncover neural mechanisms of repetitive behavior relevant to many disorders such as TS, OCD and ASD. Further, understanding mechanisms that prevent these behaviors, which may be occurring in non-stereotypy mice, has relevance for combating stereotypy behavior in brain disorders.

## Acknowledgments

This work was funded by Tourette Syndrome Association and NIH R01DA038613 (MKL), R01DC013817 (RH) and Association Française du Syndrôme de Gilles de la Tourette (ME). The authors would like to thank Sunayana Mitra, Maggie S Mattern and Makeda Turner for their technical help.

## Author contributions

ME, RH and MKL designed the experiments. ME, AL, ST and TCF conducted behavioral experiments and BE provided animal support. ME and RC conducted cell-type specific RNA extraction. ME, YS and RH performed bioinformatic analyses. ME and MEF conducted neuronal morphology analysis. ME and MKL wrote the manuscript with contributions from all authors.

The authors declare that they have no conflict of interest.

