## Supplementary Material for "Individual differences in stereotypy and neuron subtype translatome with TrkB deletion"

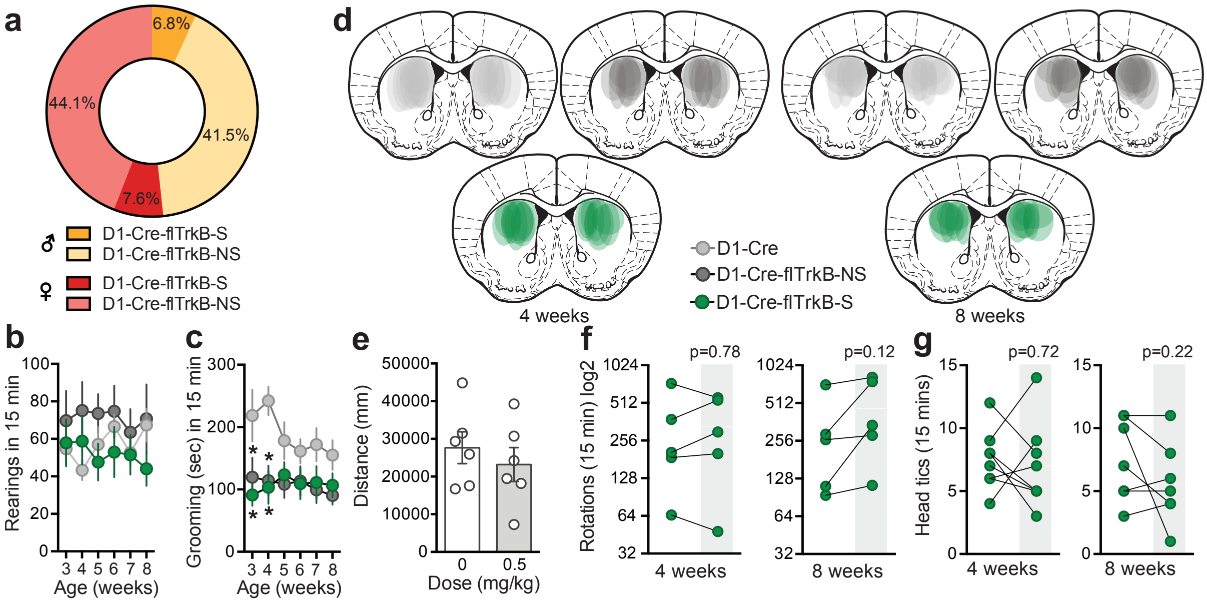


**Supplementary Figure S1:** Behavioral characterization, Clozapine-N-oxide (CNO) dose validation and control experiment for D1-MSN inhibition: **a.** Proportion of males and females in both D1-Cre-flTrkB-S and -NS groups; **b.** Rearing behavior measured from juvenile to young adult ages; **c.** Grooming behavior is lower in both D1-Cre-flTrkB-S and -NS at 3 and 4 weeks but normalizes in later ages (*p<0.05); n= 13 D1-Cre-flTrkB-S, 7 D1-Cre-flTrkB-NS, and 12 D1-Cre; **d.** AAV-DIO-hM4D(Gi)-mCherry virus placement in 4 and 8 week-old mice; **e.** No effect of the 0.5 mg/kg (ip.) CNO dose on general locomotion as measured in open field in D1-Cre mice: t-test: t= 0.73, df= 10; p= 0.49; **f.** The 0.5 mg/kg CNO dose was insufficient to impact rotational behavior in the absence of the hM4D(Gi) receptor: paired t-test: t= 0.29, df= 4; p= 0.78 and t= 2.01, df= 4; p= 0.12 at 4 and 8 weeks of age respectively; **g.** No effect of D1-MSN inhibition on head tics at 4 weeks: paired t-test: t=0.37, df=7; p= 0.72 (n= 8 in each group) and 8 weeks: t=0.37, df=6; p= 0.22 respectively (n= 8, 8, 7 respectively).

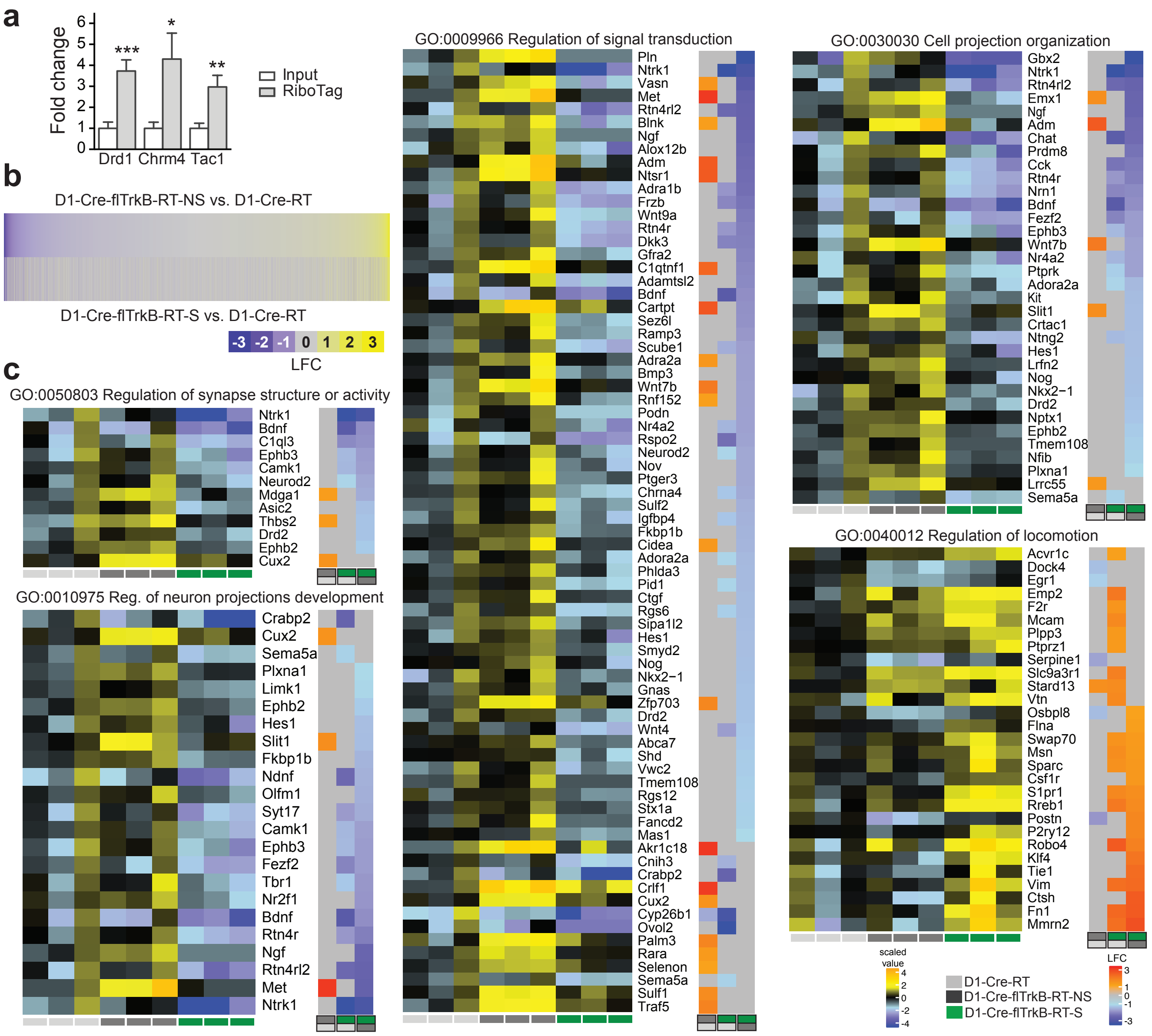


**Supplementary Figure S2**: RiboTag validation, Heat map for RNA-sequencing raw data and significant GO terms: **a.** Dorsal striatum samples display enrichment of ribosome-associated mRNA for known D1-MSN subtype genes (Input n=9, RiboTag n =11); **b.**14,764 genes were identified in D1-MSNs using RNA-sequencing; **c.** Heat maps for significant GO terms based on cluster analysis related to cell morphology and function as well as behavioral specificities: ‘Regulation of synapse structure or activity’ (GO:0050803), ‘Regulation of neuron projections development’ (GO:0010975), ‘Regulation of signal transduction’ (GO:0009966), ‘Cell projection organization’ (GO:0030030) and ‘Regulation of locomotion’ (GO:004001227). LFC= Log Fold Change. Light grey, dark grey and green labels below the heat maps indicate groups and group comparisons. * p<0.05, ** p<0.01, *** p<0.001.

**Supplementary Table 1**: Primer sequences used for multiplexed mRNA analysis with NanoString.

**Supplementary Table 2**: Raw data from D1-MSN-specific RNA-sequencing displaying read counts, Log Fold Change and False Discovery Rate.
https://drive.google.com/file/d/193EoGyvnu-hNpYDSQQOQp8uRFVb53RHG/view?usp=sharing

**Supplementary Table 3**: Post filtrated RNA-sequencing data showing significantly different expression and corresponding CLICK clusters.
https://drive.google.com/file/d/1w-AFgRy6VsB-WGQGIQVLc2qvmI3WhNOx/view?usp=sharing

**Supplementary Table 4**: Statistics for NanoString multiplexed mRNA analysis.

**Supplementary Video 1**: Representative videos showing spontaneous rotational in D1-Cre-flTrkB mouse with stereotypy compared to a D1-Cre-flTrkB mouse without stereotypy. In the D1-Cre-flTrkB-S mouse, note the head tics between bouts of rotational behaviors.
https://drive.google.com/file/d/15wYSslzL5WGgYwUGyaUv8IuDmMdI0yBp/view?usp=sharing

### TaqMan gene expression assay

| Gene name | Reference # |
|---|---|
| Gapdh | Mm99999915_g1 |
| Drd1 | Mm02620146_s1 |
| Chrm4 | Mm00432514_s1 |
| Tac1 | Mm01166996_m1 |

### NanoString primer sequences

| Gene name | Accession number | Forward sequence | Reverse sequence |
|---|---|---|---|
| Aars | NM_146217.4 | ACCCAGAAATGCATCCGA | CGCCTCATCACCACCAA |
| Adora2a | NM_009630.2 | CCCACCAAGGAATTGCAG | AAAGGCAGTAGGCAGGGC |
| Bdnf | NM_007540.4 | ACGGCTCCTGCCAATTTA | TGCTCAGGAACCCAGGAG |
| Camk1 | NM_133926.2 | AAGGCAGCATGGAGAACG | TTGACAGCATCCAGCACC |
| Chrm4 | NM_007699.2 | TGCCATCTTGTTCTGGCA | CGATGCTTGTGAACACGG |
| Dlg4 | NM_001109752.1 | TTCATTCCCAGCAAACGG | TCCAAACTTGTCGGGGAA |
| Drd1 | NM_010076.3 | CCTGAAGAGGGAGGAGGC | TCCAAAAGCAGCAGAGGG |
| Drd2 | NM_010077.2 | GCCATTGTCTGGGTCCTG | GGCTGCTACGCTTGGTGT |
| Ephb3 | NM_010143.1 | CAAGACGCTGAAGGTGGG | CCCACAAGCTGGATGACC |
| Fezf2 | NM_080433.3 | ACATGCACACCCACAACG | AGGGCAGGAAGAACGGAC |
| Gabra3 | NM_008067.4 | GCACTGTTGACTGGGGCT | AGCATCAATTGTCCCCCA |
| Gabrb1 | NM_008069.4 | CAACGGCAACAACAAAAGG | CCAGAGCACAGTGCAGGT |
| Hes1 | NM_008235.2 | TGCCTTTCTCATCCCCAA | GGGTAGCAGTGGCCTGAG |
| Hsp90ab1 | NM_008302.3 | TCCGCAAGAACATCGTCA | CTGGGTCTCCTTCATGCG |
| Klf4 | NM_010637.3 | AACTTGAACATGCCCGGA | GCTTCCTCTTCCTCCGACA |
| Limk1 | NM_010717.2 | TCTGTCTCCATCGACCCC | AGTCGATCTCGTCCAGCG |
| Macf1 | NM_001199136.1 | AGTGGGGAATCAGGGGAT | AGGCCTGCTCCAGGTTCT |
| Mcam | NM_023061.2 | ACGGCTACCCCATTCCTC | GATTCCTTCATGTGGTTCCC |
| Mme | NM_008604.3 | GCAACCTATGATGATGGCA | TTGGGTTCTTGAAGGACATCTT |
| Ndnf | NM_172399.3 | GTGGGTGCCTTTGTCAGG | GACAGAAGCAGCCTCCCA |
| Neurod2 | NM_010895.3 | ACAGGCCAGAGGCAGTTG | ATCCACATCCCCTCCCAC |
| Nr2f1 | NM_010151.2 | GATTGCCCTCTCTTCGCA | AGGGAGTCAGGGAGCAGG |
| Ntrk1 | NM_001033124.1 | ACTGAGGGCAAAGGCTCC | TCAGTGCCTTGACAGCCA |
| Ntrk2 | NM_001025074.1 | AAGCCACACACAGGGCTC | GAACCACTGAAGCGCAGG |
| Pdyn | NM_018863.3 | CGTCCTGAAGGAGCTGGA | CGGAACTCCTCTTGGGGT |
| Penk | NM_001002927.2 | CATGAGAAGGGTGGGACG | AGCAGGCTAGTGGGGGTC |
| Pum1 | NM_001159605.1 | TGGAGGAGCTAGCCAACG | AGCTCCGAGGCCATTTCT |
| Rreb1 | NM_001177868.1 | TCACGATGCCGAGTCAGA | CAGTCCTCGATGTGTGCG |
| Rtn4rl2 | NM_199223.1 | TGACCCCCAGCTGTCCTA | TGGAGAAGAGCCACAGGG |
| S1pr1 | NM_007901.4 | ACCTGCTGTTGTCTGGGG | CCCCAGGATGAGGGAGAT |

| | | | |
|---|---|---|---|
| Scn1b | NM_011322.2 | CTGCGTGGAGGTGGATTC | CCCGACTACCGTTCCACA |
| Serpine1 | NM_008871.2 | AAATGTCCACCTTGCCCA | ACGAAGAGCCAGGCACAC |
| Shank1 | NM_001034115.1 | AGCACCGACCCAAAGGAT | AGTGGGTGGTGGGGGTAT |
| Slc35d3 | NM_029529.3 | GTGCTGGGCATCTCGGT | CAGCGAAGGAGCGAGCTA |
| Smyd2 | NM_026796.1 | TACAAAGGGACCCTGGCA | GCTGCTGAGCTTTCGGAC |
| Sparc | NM_009242.4 | AGGAGGAGGAGGGCCTTT | AACTCTCGGCACAGGCAG |
| Strn | NM_011500.2 | TGAGCAGTGCTACAGCGG | CGAAGAGTGCCGTCTGCT |
| Syt17 | NM_138649.1 | GCTTCAAAGTTCCCCAGGA | GCAGTGCGGTGAGTGTTG |
| Tac1 | XM_006505028.1 | CAGAGAATCGCCCGAAGA | TGCGTTCAGGGGTTTATTT |
| Tbr1 | NM_009322.3 | CACAGCTAGGCCGCCA | CAAGGTCGGTGCGCTAAC |
| Vim | NM_011701.4 | TCTGCCACTCTTGCTCCG | GAGCCACCGAACATCCTG |
| Vtn | NM_011707.2 | CTCTGCCCAAGCCAAAAA | GACGCTCTGAATGGGCTC |
| Wnt4 | NM_009523.1 | ACCATGAGCCCCCGTT | ATCTGCACCTGCCTCTGG |
| Wnt9a | NM_139298.2 | AAGACCCAGACTTGGGGG | AGGGGGTACACTCCTGCC |

Engeln et al. Suppl Table 4

| Gene name | Accession # | 4-week old ANOVA | 4-week old Post hoc | 8-week old ANOVA | 8-week old Post hoc |
|---|---|---|---|---|---|
| Adora2a | NM_009630.2 | p=0.17 | p=0.14 (NS vs. S) | F (2, 19) = 17.01; p<0.0001 | p<0.01 (Ctrl vs. NS and NS vs. S) |
| Bdnf | NM_007540.4 | p=0.18 | p=0.2 (Ctrl vs. S) | F (2, 18) = 3.556; p=0.05 | p<0.05 (NS vs. S) |
| Camk1 | NM_133926.2 | p=0.69 | n.s. | p=0.52 | n.s. |
| Chrm4 | NM_007699.2 | F (2, 17) = 4.354; p<0.05 | p<0.05 (NS vs. S) | p=0.3 | n.s. |
| Dlg4 | NM_001109752.1 | p=0.88 | n.s. | p=0.91 | n.s. |
| Drd1 | NM_010076.3 | p=0.4 | n.s. | p=0.21 | n.s. |
| Drd2 | NM_010077.2 | p=0.76 | n.s. | F (2, 19) = 22.17; p<0.0001 | p<0.001 (Ctrl vs. NS and NS vs. S) |
| Ephb3 | NM_010143.1 | p=0.43 | n.s. | p=0.11 | p<0.18 (Ctrl vs. S and NS vs. S) |
| Fezf2 | NM_080433.3 | p=0.37 | n.s. | K-W= 6.64; p<0.05 | p=0.05 (NS vs. S) |
| Gabra3 | NM_008067.4 | p=0.52 | n.s. | F (2, 19) = 3.317; p=0.05 | p<0.05 (NS vs. S) |
| Gabrb1 | NM_008069.4 | p=0.17 | p=0.19 (Ctrl vs. S) | F (2, 18) = 3.930; p<0.05 | p<0.05 (Ctrl vs. S) |
| Hes1 | NM_008235.2 | p=0.34 | n.s. | p=0.11 | p=0.09 (NS vs. S) |
| Klf4 | NM_010637.3 | N.D. | N.D. | p=0.69 | n.s. |
| Limk1 | NM_010717.2 | p=0.72 | n.s. | F (2, 19) = 2,756; p=0.08 | p=0.09 (NS vs. S) |
| Macf1 | NM_001199136.1 | F (2, 17) = 5.656; p<0.05 | p<0.05 (Ctrl vs. S and NS vs. S) | p=0.24 | n.s. |
| Mcam | NM_023061.2 | N.D. | N.D. | F (2, 18) = 3.084; p=0.07 | p=0.06 (Ctrl vs. NS) |
| Mme | NM_008604.3 | F (2, 17) = 5.510; p<0.05 | p<0.05 (Ctrl vs. S and NS vs. S) | F (2, 19) = 6.160; p<0.01 | p<0.01 (Ctrl vs. S) |
| Ndnf | NM_172399.3 | K-W = 5.757; p<0.05 | p=0.05 (Ctrl vs. S) | F (2, 19) = 8.832; p<0.01 | p<0.05 (Ctrl vs. NS and NS vs. S) |
| Neurod2 | NM_010895.3 | F (2, 15) = 3,161; p=0.07 | p=0.06 (Ctrl vs. NS) | p=0.12 | p<0.19 (Ctrl vs. S and NS vs. S) |
| Nr2f1 | NM_010151.2 | p=0.27 | n.s. | K-W= 7.524; p=0.02 | p<0.05 (NS vs. S) |
| Ntrk1 | NM_001033124.1 | p=0.27 | n.s. | F (2, 19) = 13.39; p<0.001 | p<0.01 (Ctrl vs. NS and NS vs. S) |
| Ntrk2 | NM_001025074.1 | F (2, 16) = 54.68; p<0.0001 | p<0.0001 (Ctrl vs. NS and S) | F (2, 18) = 62,94; p<0.0001 | p<0.0001 (Ctrl vs. NS and S) |
| Pdyn | NM_018863.3 | p=0.69 | n.s. | F (2, 19) = 8.705; p<0.01 | p<0.05 (Ctrl vs. NS and NS vs. S) |
| Penk | NM_001002927.2 | F (2, 16) = 2,957; p=0.08 | p=0.07 (Ctrl vs. S) | F (2, 19) = 15,48; p<0.0001 | p<0.01 (Ctrl vs. S and NS vs. S) |
| Rreb1 | NM_001177868.1 | p=0.3 | n.s. | p=0.86 | n.s. |
| Rtn4rl2 | NM_199223.1 | p=0.24 | n.s. | F (2, 17) = 3.320; p=0.06 | p=0.05 (NS vs. S) |
| S1pr1 | NM_007901.4 | F (2, 17) = 6.517; p<0.01 | p<0.05 (Ctrl vs. S and NS vs. S) | F (2, 19) = 3.267; p=0.06 | p=0.05 (Ctrl vs. NS) |
| Scn1b | NM_011322.2 | p=0.53 | n.s. | F (2, 19) = 5,384; p<0.05 | p<0.05 (Ctrl vs. S) |
| Serpine1 | NM_008871.2 | N.D. | N.D. | F (2, 19) = 7.751; p<0.01 | p<0.05 (Ctrl vs. NS and S) |
| Shank1 | NM_001034115.1 | p=0.8 | n.s. | F (2, 18) = 10.34; p<0.01 | p<0.01 (Ctrl vs. S and NS vs. S) |
| Slc35d3 | NM_029529.3 | p=0.49 | n.s. | p=0.5 | n.s. |
| Smyd2 | NM_026796.1 | p=0.91 | n.s. | F (2, 19) = 4.044; p<0.05 | p<0.05 (NS vs. S) |
| Sparc | NM_009242.4 | N.D. | N.D. | p=0.16 | p=0.15 (NS vs. S) |
| Strn | NM_011500.2 | F (2, 17) = 3.437; p=0.055 | p=0.09 (Ctrl vs. S and NS vs. S) | F (2, 19) = 6.548; p<0.01 | p<0.05 (Ctrl vs. S and NS vs. S) |
| Syt17 | NM_138649.1 | p=0.92 | n.s. | F (2, 18) = 5.144; p<0.05 | p<0.05 (NS vs. S) |
| Tac1 | XM_006505028.1 | p=0.51 | n.s. | p=0.69 | n.s. |
| Tbr1 | NM_009322.3 | p=0.74 | n.s. | F (2, 18) = 3.215; p=0.06 | p=0.06 (NS vs. S) |
| Vim | NM_011701.4 | p=0.15 | p=0.18 (NS vs. S) | p=0.23 | n.s. |
| Vtn | NM_011707.2 | p=0.22 | n.s. | p=0.67 | n.s. |
| Wnt4 | NM_009523.1 | N.D. | N.D. | F (2, 18) = 16.29; p<0.0001 | p<0.05 (Ctrl vs. NS and S) |
| Wnt9a | NM_139298.2 | p=0.83 | n.s. | F (2, 18) = 7.545; p<0.01 | p<0.05 (Ctrl vs. NS and NS vs. S) |

p<0.05 or less

p≤0.09

Not detected N.D.
